# Trade-off between plasticity and velocity in mycelial growth

**DOI:** 10.1101/2020.10.25.354373

**Authors:** Sayumi Fukuda, Riho Yamamoto, Naoki Yanagisawa, Naoki Takaya, Yoshikatsu Sato, Meritxell Riquelme, Norio Takeshita

## Abstract

Tip-growing fungal cells maintain the cell polarity at the apical regions and elongate by de novo synthesis of cell wall. Cell polarity and growth rate affect the mycelial morphogenesis, however, it remains unclear how they act cooperatively to determine cell shape. Here we investigated their relationship by analyzing hyphal tip growth of filamentous fungi growing inside extremely narrow 1 μm-width channels of microfluidic devices. Since the channels are much narrower than the diameter of hyphae, the hyphae must change its morphology when they grow through the channels. Live imaging analysis revealed that hyphae of some species continued growing through the channels, whereas hyphae of other species often ceased growing when passing through the channels or lost the cell polarity after emerging from the channels. Fluorescence live imaging analysis of the Spitzenkörper, a collection of secretory vesicles and polarity-related proteins at hyphal tips, in *Neurospora crassa* hyphae indicates that hyphal tip growth requires a very delicate balance of ordered exocytosis to maintain polarity in spatially confined environments. We analyzed the mycelial growth of seven fungal species from different lineages, which also include phytopathogenic fungi. This comparative cell biology showed that the growth defects in the channels were not correlated with their taxonomic classification nor with the width of hyphae, but, correlated with the hyphal elongation rate. This is the first report indicating a trade-off between plasticity and velocity in mycelial growth, and serves to understand fungal invasive growth into substrates or plant/animal cells, with direct impact on fungal biotechnology, ecology and pathogenicity.

## Introduction

Cell morphogenesis, which is controlled by cell polarity and cell growth, is fundamental for all cellular functions (1,2). The core cell polarity machinery appears to be relatively conserved in animals, plants, and fungi (3, 4). First, polarity signaling complexes assemble near a cell-surface landmark, and locally assemble the cytoskeleton through actin or tubulin polymerization. Then, directed trafficking of vesicles and carriers contribute to local membrane and cell wall expansion. In addition, cell growth is controlled by turgor pressure, which drives the expansion of the cell membrane, especially in cell types covered by a cell wall (5,6). Although both polarity and growth are essential for cell morphogenesis, how growth speed and cell polarity cooperatively control cell shape remains unclear.

Filamentous fungi grow as highly polarized tubular cells by elongation of their primary hyphae and branches at the tips (7). The tip-growing fungal cells maintain polarity at the apical regions, where they elongate by supply of membrane lipids and de novo synthesis of cell wall (8–11). The necessary proteins and lipids are delivered to the tip by vesicle trafficking via the actin and microtubule cytoskeletons and their corresponding motor proteins (12–16). The delivered secretory vesicles accumulate temporarily in an apical vesicle cluster, called the Spitzenkörper (SPK; 17–19). Vesicle exocytosis at the apical membrane allows release of secretory enzymes and the expansion of apical membrane and cell wall. Recent live imaging analyses including super-resolution microscopy have revealed that the multiple steps in polarized growth, such as the assembly of polarity markers, actin polymerization, and exocytosis, are temporally coordinated through pulsed Ca^2+^ influxes (20–22).

While tip growth rate depends on the supply of vesicles, it has been reported that turgor pressure is one of the major forces driving the expansion at the hyphal tip (6). Turgor pressure in growing hyphae has been directly measured by using microinjection with pressure probe (23). Cytoplasmic bulk flow, which is evident in fast growing fungi like *Neurospora crassa*, is also involved in the force to expand the hyphal tip (6, 24).

Microfluidic devices-based technology has been used to study the behavior of tip-growing plant cells (25–27) and, more recently, of filamentous fungi (28, 29). An elastic polydimethylsiloxane (PDMS) microfluidic device enabled to measure the invasive pressure of tip-growing plant pollen tubes (30). Likewise, scanning probe microscopy (SPM), with a sensor probe that directly indents the cellular surface, is available for measurement of cellular stiffness in a non-invasive manner (31). These methods in combination with cell biology are powerful tools to investigate the mechanical properties in living cells.

Here we constructed a microfluidic device with 1 μm-width channels, which are narrower than the diameter of fungal hyphae, and observed growth as hyphae grew into, through and out of the channels (Fig. 1A). The present study aimed to identify the relationship between cell polarity and growth rate by observing the forced morphological changes of growing hyphae under a microscope. Our results would help to understand fungal invasive growth into substrates or host plant/animal cells, and apply that knowledge to the fields of fungal biotechnology, ecology and pathogenicity.

**Fig. 1.**
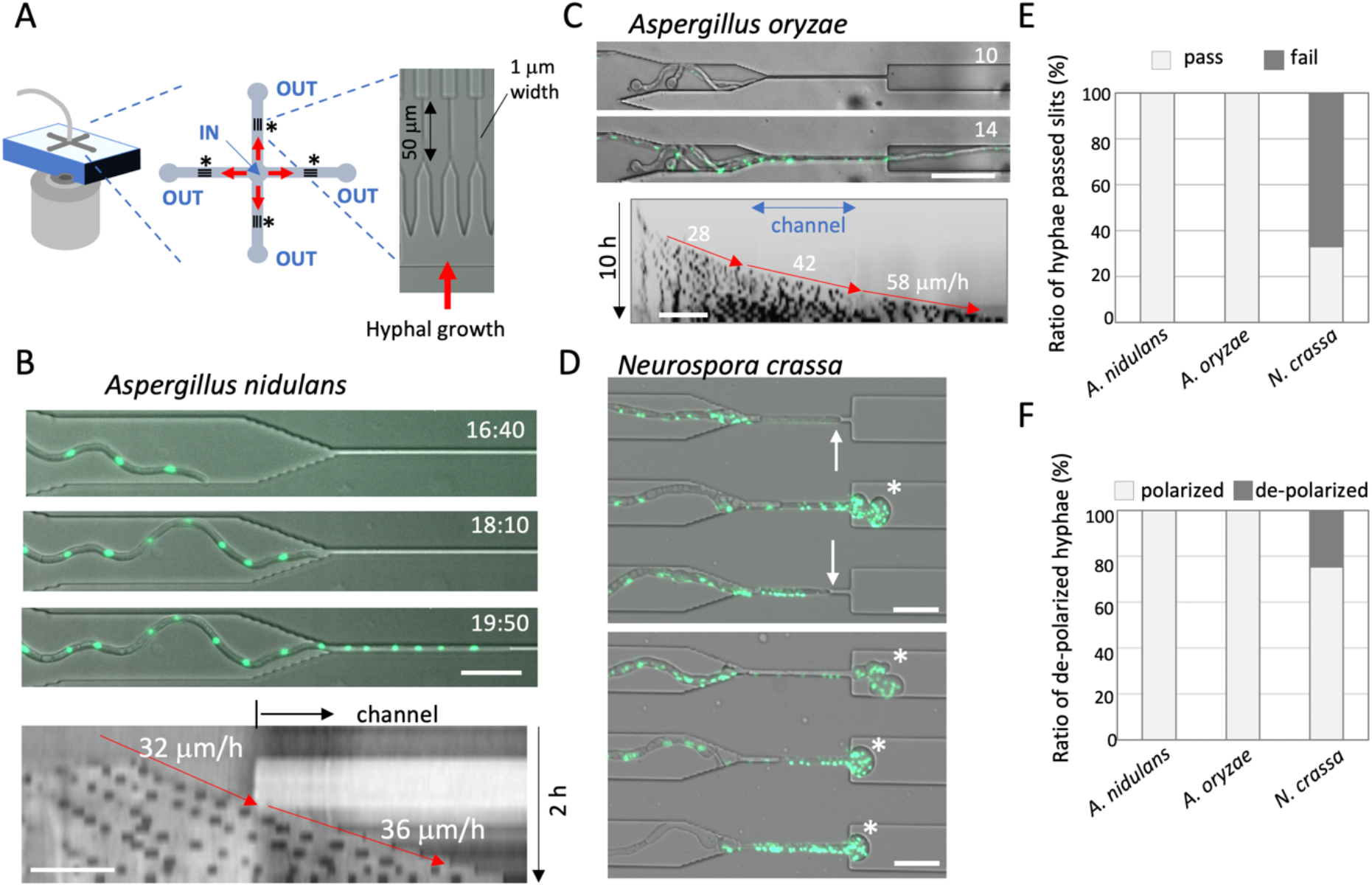
*A. nidulans* and *A. oryzae* but not *N. crassa* hyphae passed through the channels. (A) A design of microfluidic device; Inflow at the center “IN” and outflows at the four path ends “OUT”. Twenty channels of 1 μm width and 50 μm length were designed between the inlet and outlet per one path (asterisks). (B) Time series showing a hypha of *A. nidulans* (nuclei labeled with GFP) growing into the channel. The elapsed time is given in hours:minutes. Kymograph along the growth axis before and in the channel from Movie S1. The hyphal elongation rates before and in the channel are shown by arrows. Total 2 h, scale bar: 20 μm. (C) Time series showing a hypha of *A. oryzae* (nuclei labeled with GFP) hypha passed through the channel. The elapsed time is given in hours. Kymograph along the growth axis before, in and after the channel from Movie S2. Total 10 h, scale bar: 20 μm. (D) Images of *N. crassa* (nuclei labeled with GFP) hyphae stopped growing in the channels (arrows) and de-polarized hyphae exiting from the channels (asterisks) from Movie S3, scale bar: 20 μm. (E) Ratio of the hyphae that successfully passed through the channels (pass) or stopped in or exiting from the channels (fail) in *A. nidulans*, *A. oryzae* and *N. crassa.* n=50, respectively. (F) Ratio of polarized or de-polarized hyphae that passed through the channels in *A. nidulans*, *A. oryzae* and *N. crassa.* n=50, respectively.

## Results

### *A. nidulans* and *A. oryzae* but not *N. crassa* hyphae grow through the channels

The PDMS microfluidic device used in this study possesses multiple micro channels, 1 μm wide and 50 or 100 μm long (Fig. 1A). Fungal spores were inoculated at the center of the device, “IN”. The medium solution was continuously supplied to the inlet “IN” with the help of a pump (0.8 μl per hour) and flowed out from the four outlet “OUT” corners.

We monitored the hyphal growth of *Aspergillus nidulans* as it grew into, through and out of the channels. We used a strain whose nuclei were visualized by GFP, nuclear localization signal of the transcription factor StuA tagged with GFP (32). The hyphal widths were 2-3 μm before entering the channel under this condition. All observed hyphae grew into the channels, passed through them and continued to grow (50<n) (Fig. 1B, S1A, Movie S1). The kymograph along the growth axis indicated comparable growth rate, 37 ± 15 μm/h (n=20), before, through (Fig. 1B) and after the channels (Fig. S1A). In some cases, two or three hyphae passed through the same channel (Fig. S1B, Movie S1). In the same way, we tested *Aspergillus oryzae*, which is important for traditional food fermentation and modern biotechnology (33). We used a strain in which histone H2B is fused with GFP (34). Again, all observed hyphae went into the channels, passed through there and continued to grow without growth rate decrease (84 ± 37 μm/h, n=30) (Fig. 1C, Movie S2).

We examined another model filamentous fungus, *Neurospora crassa*, whose hyphae usually grow faster and have a larger diameter than those in *A. nidulans* (7, 35)(see below). We used a strain in which histone H1 is fused with GFP (36). Some hyphae penetrated into the channels but often their growth speed slowed down and stopped before reaching the end of the channel (Fig. 1D arrows, Movie S3). The hyphae that passed through channels frequently lost polarized growth and started to swell (Fig. 1D asterisks, Movie S3). The de-polarized hyphae stopped growing after a while, then lost the GFP signal of nuclei (Movie S3). The growth arrest inside the channels and the loss of polarity of the hyphae after exiting the channels were characteristic of *N. crassa* but not of *A. nidulans* or *A. oryzae* (Fig. 1E, F). A 33 % of *N. crassa* hyphae grew out of the channel without losing the cell polarity (Fig. 1E, n=50). In addition, the *N. crassa* spores that were trapped in front of the channels frequently germinated to the opposite side of channels (Fig. S1C), but not in the case of *A. nidulans* or *A. oryzae* (Fig. S1D).

### Cell polarity loss after forced morphological changes in *N. crassa*

We investigated the cell polarity in *N. crassa* hyphae growing in the channels by monitoring GFP tagged CHS-1 (chitin synthase class III) at the SPK (37). Accumulation of GFP-CHS-1 at the SPK was clearly observed at the tips of growing hyphae before growing into the channels (Fig. 2A, Fig. S1E, Movie S4). The hyphae penetrated into the channels then stopped growing, coinciding with a loss of the GFP signal at the SPK, and dispersion of the fluorescence signal along the cytoplasm of the tip region with high intensity level of GFP (Fig. 2A, B, Fig. S1D). Distinct accumulation of GFP-CHS-1 at the SPK was hardly observed in the hyphae growing within the channels (Fig. 2C). In the de-polarized swollen hyphae exiting the channel, the fluorescence signal was diffused and weak, but became visible again at the SPK of the multiple branches that formed when polarized growth resumed (Fig. 2D arrows, Movie S5). The kymograph along the growth axis indicated comparable growth rate before and in the channels, 200 and 245 μm/h, however the hypha drastically decreased the growth rate after exiting the channel (Fig. 2E).

**Fig. 2.**
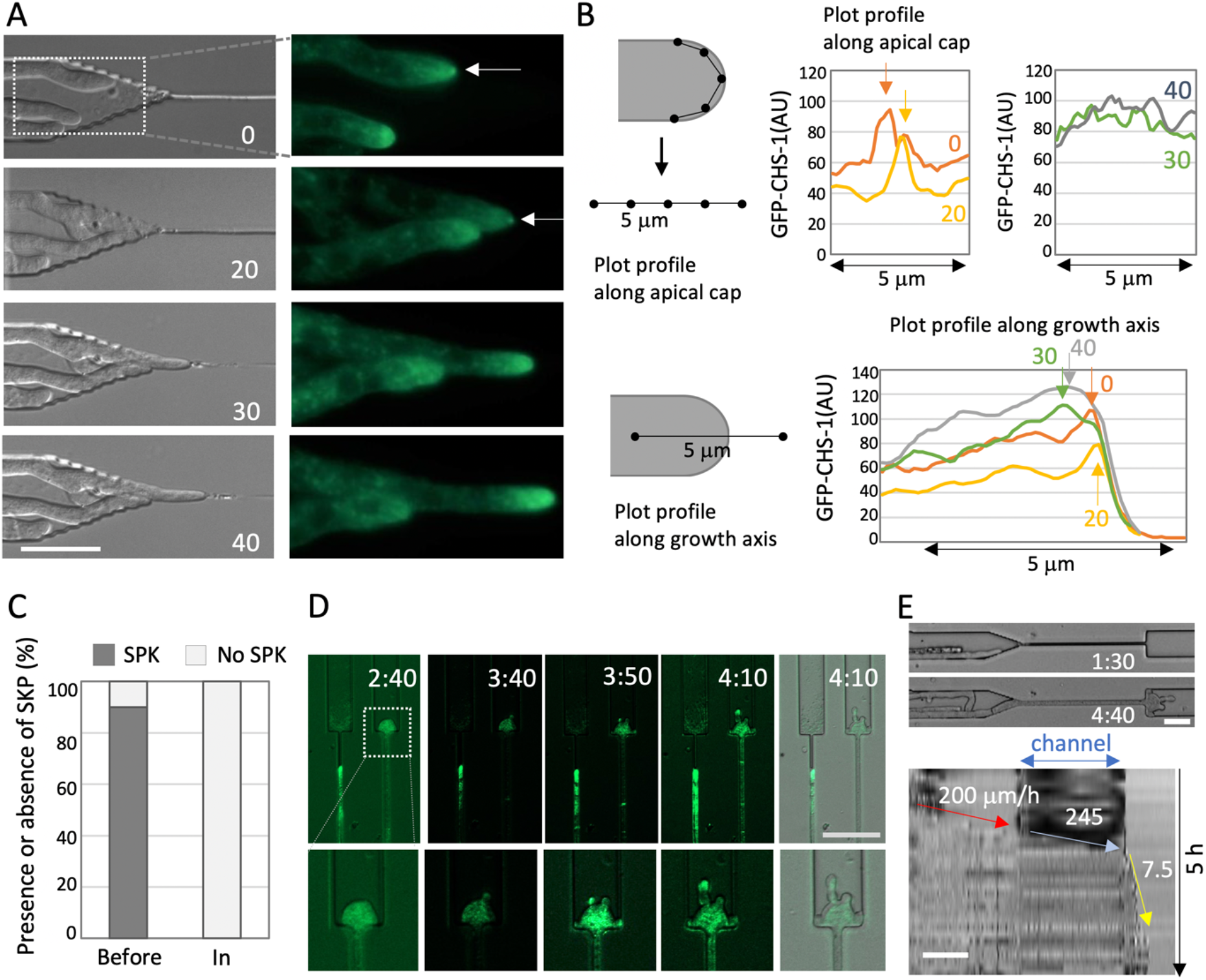
Cell polarity and septum formation during confined growth in *N. crassa*. (A) Time series images of *N. crassa* (DIC; left, CHS-1-GFP; right) hyphae growing into a channel from Movie S4. The arrows indicate the SPK. The elapsed time is given in minutes. Scale bar: 20 μm. (B) Scheme to measure GFP signal intensity along the apical membrane (upper) or the growth axis (lower). The plot profile along the apical membrane (upper) indicates the signal intensity peaks of the SPK (arrows) at 0, 20 min, but not at 30, 40 min. The plot profile along the growth axis (lower) indicates the peaks at the apex of hyphae at 0, 20 min, but at the sub-apex at 30, 40 min. (C) Ratio of presence or absence of SPK in hyphae before or in channels. n=20, 10. (D) Image sequence of the de-polarized hypha after exiting the channel in the *N. crassa* (CHS-1-GFP) hypha from Movie S5. The elapsed time is given in hours:minutes. Scale bar: 50 μm. (E) Kymographs along the growth axis of the channel from Movie S5. The hyphal elongation rates before entering, through and after exiting the channel are shown by arrows. Total 5 h, scale bar: 50 μm.

CHS-1-GFP is also known to localize at septa during their formation (37). We found that the de-polarized hyphae possessed two-times more septa within the narrow channels than the hyphae that successfully passed through the channels without presenting polarity defects (Fig. S2A-C), suggesting that the cell cycle progresses and deposition of cross walls continues when tip growth is inhibited.

### Relationship between growth rate and polarity maintenance

We tested two plant pathogenic fungi, *Fusarium oxysporum* and *Colletotrichum orbiculare* (38, 39), using the same microfluidic devices. Since the plant pathogenic fungal hyphae have to invade between tightly connected plant cells, polarity maintenance in spatially confined growth should be important for their pathogenicity. Almost all the hyphae of *F. oxysporum* and *C. orbiculare* grew into and passed through the channels while maintaining the growth rate (83 ± 27 μm/h and 91 ± 16 μm/h, n=53, 45, respectively) (Fig. 3A, Movie S6, 7).

**Fig. 3.**
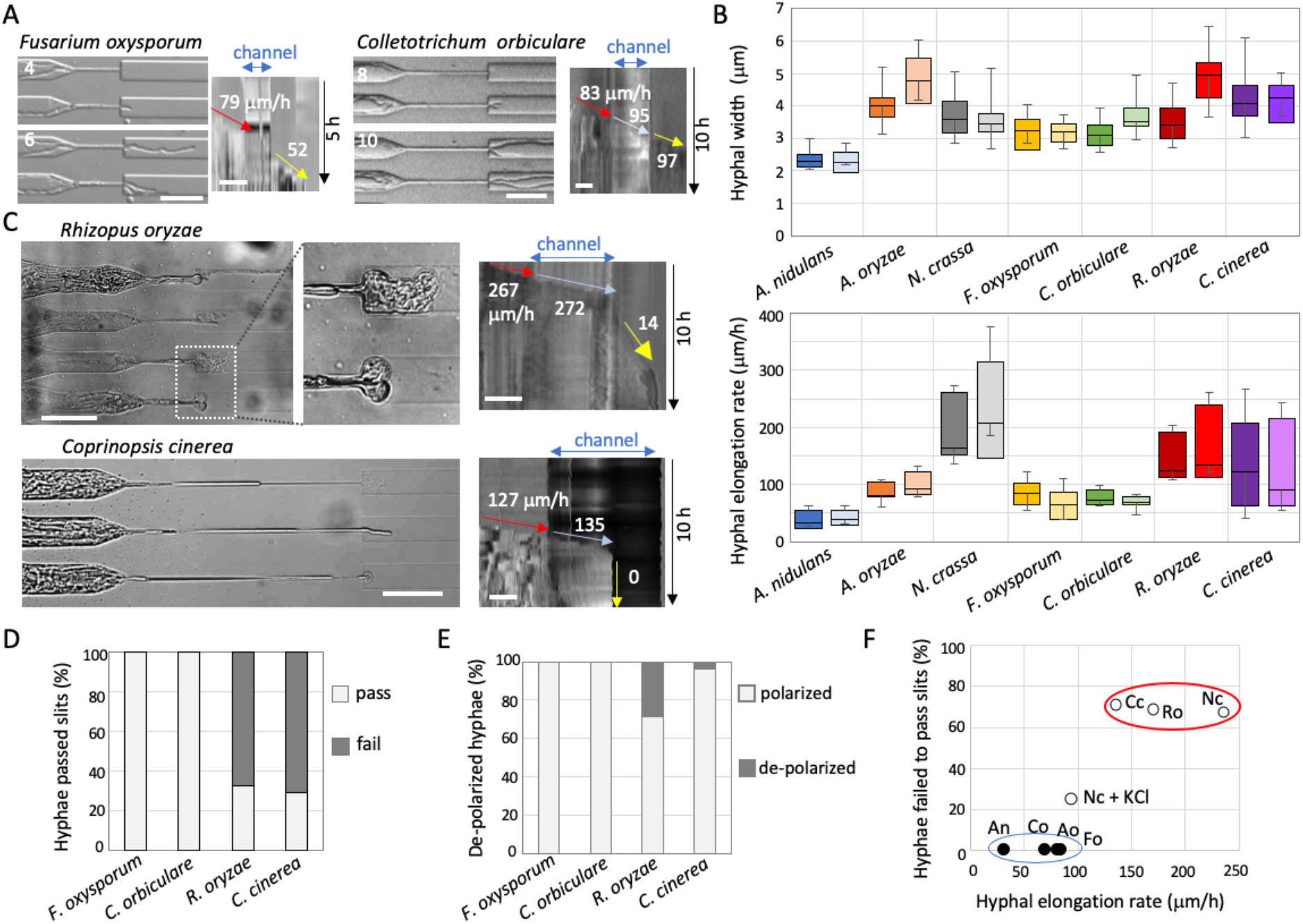
Relationships among hyphal width, growth rate and polarity maintenance. (A) Time series of *F. oxysporum* (left) and *C. orbiculare* (right) hyphae that passed through the channel (from Movies S6, 7). The elapsed time is given in hours. Kymographs along the growth axis of the channel from Movie S10 and S11. The hyphal elongation rates before entering and after exiting the channel are shown by arrows. Total 5 h (left) and 10 h (right), scale bar: 50 μm. (B) Boxplots of hyphal width (upper) and hyphal elongation rate (lower) in *A. nidulans, A. oryzae, N. crassa, F. oxysporum, C. orbiculare, R. oryzae* and *C. cinerea* before entering the channels (dark color) and after exiting the channels (light color). n=26, 40, 80, 53, 45, 14, 20, 80 (upper), n=20 (lower). De-polarized hyphae were not counted. (C) Images of de-polarized hyphae of *R. oryzae* (upper) and of *C. cinerea* hyphae that stopped growing in the channel or after exiting the channel (lower), from Movies S8, 9. Kymograph along the growth axis of the channel. Total 10 h, scale bars: 50 μm. Scale bars: 20 μm. (D, E) Ratio of the hyphae that successfully passed through the channel (pass) or stopped in or just after exiting the channels (fail) (D), and ratio of de-polarized hyphae after exiting the channels (E) in *F. oxysporum, C. orbiculare, R. oryzae* and *C. cinerea*. n=29, 20, 52, 52, respectively. (F) Correlation between the hyphal elongation rate with the growth defect in channels. Two groups are shown by red or blue ellipses.

To investigate the reason why only *N. crassa* but not the other fungi showed growth defects in spatially confined growth that they were subjected to in the channels, we compared the widths of hyphae and hyphal elongation rates of all fungi grown in the device (Fig. 3B). The results corresponding to before entering and after exiting the channels are shown in dark and bright colors, respectively. The hyphal widths in *A. nidulans* were 2-3 μm, whereas those in *N. crassa, F. oxysporum* and *C. orbiculare* were 3-4 μm, and those in *A. oryzae* were slightly wider. These results suggest that the hyphal widths are not correlated to the growth defect shown in the channels. There is no significant difference in the the hyphal widths between before entering and after exiting the channels except *A. oryzae*. It is known that *A. oryzae* increases hyphal width as cultivation time passes (34). Since the widths in mature hyphae of *N. crassa* are known to be over 10 μm, the hyphae we observed in this condition were considered as young hyphae.

In contrast, the hyphal elongation rate in *A. nidulans* was less than 50 μm/h, whereas those in *A. oryzae, F. oxysporum* and *C. orbiculare* were 50-100 μm/h (Fig. 3B, lower graph). Notably, the hyphal elongation rate in *N. crassa* was 150-250 μm/h, higher than that of the other fungi.

To examine the relationship between growth rate and growth defect in the channels, we tested also *Rhizopus oryzae* and *Coprinopsis cinerea* dikaryon, whose hyphal elongation rates are known to be relatively high (40, 41). The hyphal elongation rates of *R. oryzae* and *C. cinerea* in the device were 100-250 μm/h (Fig. 3B), whereas the hyphal widths were 3-5 μm, indicating certainly that these two fungi grow faster than the other fungi, and similar to *N. crassa*. At least one hypha penetrated into one channel, however the hyphae of *R. oryzae* and *C. cinerea* often stopped growing in or shortly after exiting the channels (Fig. 3C, D, Movie S8, 9). De-polarized hyphae were sometimes observed after exiting the channels in *R. oryzae* similarly to what was observed for *N. crassa* (Fig. 3C, E). We tested various fungal species of different phylogenetic lineages (40), however the observed output did not correlate with the phylogenetic distance (Fig. S3A). Altogether these results indicated that neither phylogenetic relevance nor the width of hyphae were correlated with the growth defect in the channels. In contrast, the hyphal elongation rate displayed a strong correlation with the growth defects in the channels (Fig. 3F, Fig. S3B, C).

### Contribution of turgor pressure for polarity maintenance

Why do hyphae of *N. crassa* and *R. oryzae* generally grow faster than those of *A. nidulans* and other species? One possibility points to the fact that *N. crassa* and *R. oryzae* hyphae have higher turgor pressure. This is supported by the results showing that both *N. crassa* and *R. oryzae* were sensitive to the high osmotic condition generated by addition of 0.6 M KCl, resulting in decrease of turgor pressure (Fig. 4A and Fig. S3D). In contrast, *A. nidulans, A. oryzae,* and *F. oxysporum* were not sensitive to the high osmotic condition.

**Fig. 4.**
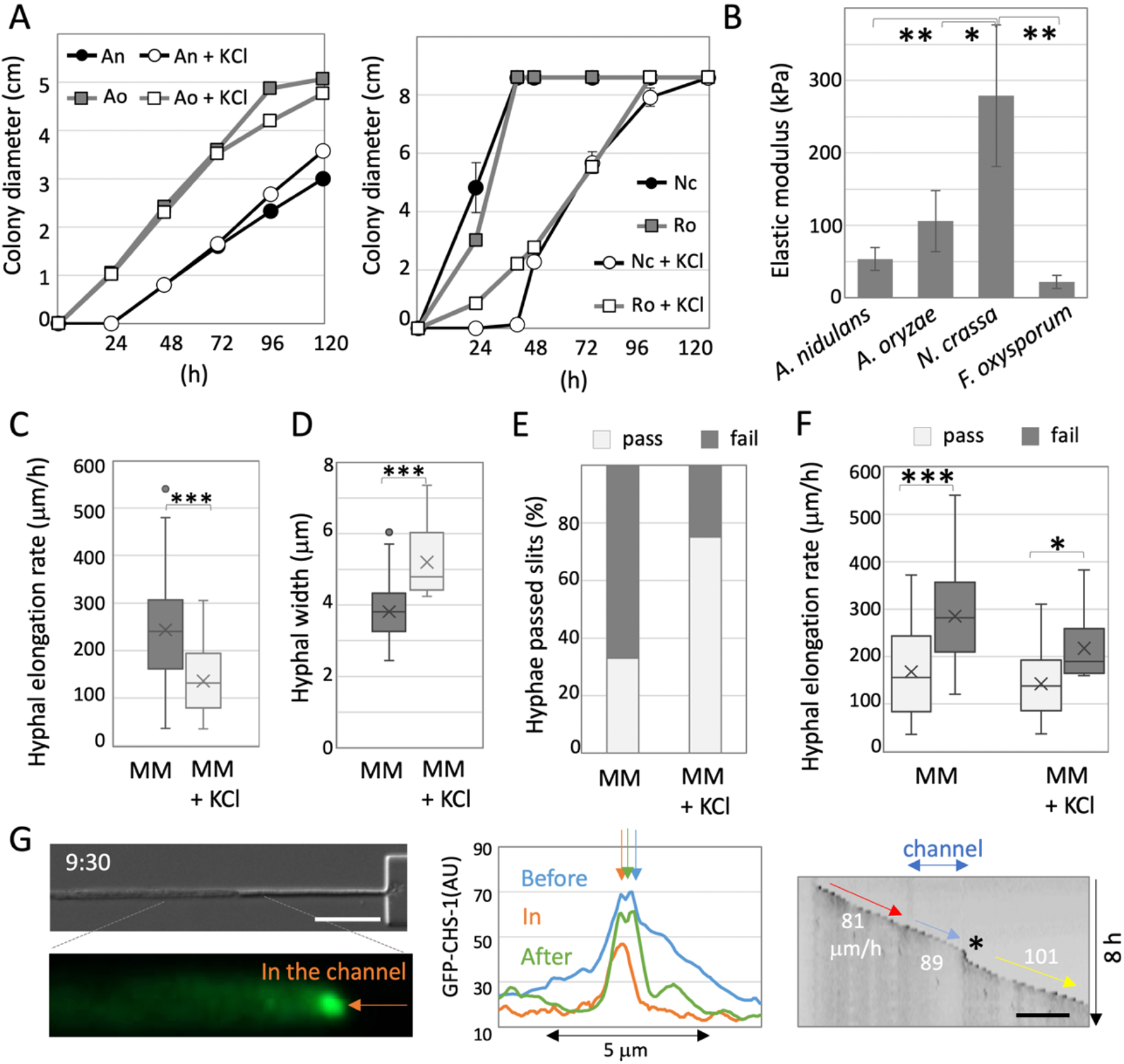
Contribution of growth rate for polarity maintenance. (A) Colony diameter of *A. nidulans* and *A. oryzae* (left)*, N. crassa* and *R. oryzae* (right) on MM or MM + 0.6 M KCl plates. (B) Elastic modulus measured by a scanning probe microscope in the hyphae of *A. nidulans, A. oryzae, N. crassa* and *F. oxysporum* grown in MM. Error bar: S.D., n = 9 in 3 hyphae, ** P ≤ 0.01, * P ≤ 0.05. (C) Hyphal elongation rate of *N. crassa* hyphae grown in MM and MM + 0.6 M KCl. Error bar: S.D., n = 20, *** P ≤ 0.001. (D) Hyphal width of *N. crassa* grown in MM and MM + 0.6 M KCl. Error bar: S.D., n = 20, *** P ≤ 0.001. (E) Ratio of the hyphae that successfully passed through the channel (pass) or stopped within or just after exiting the channels (fail) in *N. crassa* grown in MM or MM + 0.6 M KCl. n = 53, 32. (F) Boxplots of hyphal elongation rate in *N. crassa* hyphae just before entering the channel, pass or fail, grown in MM or MM + 0.6 M KCl. n = 18, 33, 18, 9. *** P ≤ 0.001. * P ≤ 0.05. (G) Image of a *N. crassa* hypha (SPK labeled with GFP) growing within the channel; from Movie S15. The arrow indicates the SPK. The elapsed time is given in hours:minutes. Scale bar: 20 μm. The plot profile along the apical membrane indicated the signal peaks of SPK (arrows) before, in and after the channel, see Fig. S4. Kymograph of GFP signal along the growth axis from Movie S10. Total 8 h, scale bar: 100μm. The hyphal elongation rates before entering and after exiting the channel are shown by arrows.

Indeed, we measured the elastic modulus, that represent forces balanced in the opposite direction of turgor pressure, by using a scanning probe microscope (SPM). The SPM scanned sample surfaces with an extremely sharp sensor probe and measured the physical property of fungal cells in a non-invasive manner at high magnifications (Fig. S4). The elastic modulus in *N. crassa* hyphae, 278 ± 98 kPa, were significantly higher than those in *A. nidulans, A. oryzae,* and *F. oxysporum* (Fig. 4B), indicating that the turgor pressure in *N. crassa* is higher than those in others.

In order to decrease the turgor pressure in hyphae of *N. crassa* grown in the device*, N. crassa* was grown in the high osmotic condition with 0.6 M KCl. The hyphal elongation rate just before the channels decreased in high osmotic condition from 239 ± 50 to 151 ± 21 μm/h (Fig. 4C), whereas the hyphal widths increased from 3.8 ± 0.7 to 5.2 ± 1.0 μm (Fig. 4D). Notably, the ratio of hyphae that passed through the channels increased from 33 to 75% in the high osmotic condition (Fig. 4E), which is correlated with the decreased hyphal elongation rate (Fig. 3F, Nc + KCl). Although 25% hyphae still stopped growing in the channels, de-polarized hyphae were not observed (Fig. S3B). We compared the hyphal elongation rate just before the channels in the hyphae that passed or failed to pass, and found that hyphal elongation rate is lower in the hyphae that passed the channels than that in the hyphae failed to pass in normal and high osmotic conditions (Fig. 4F).

Under the high osmotic condition, the SPK labelled by GFP-CHS-1 was clearly observed at the tips of growing hyphae even in the channels (Fig. 4G, Fig. S5, Movie S10). Although the hypha swelled slightly when exiting the channel (Fig. 4G right, asterisk, Movie S10), the hypha grew into and passed through the channels while maintaining the growth rate (Fig. 4G right, arrows). These results indicate that the growth rate is important for the maintenance of cell polarity in spatially confined growth derived from passing squeezed through the channels (Fig. 5).

**Fig. 5.**
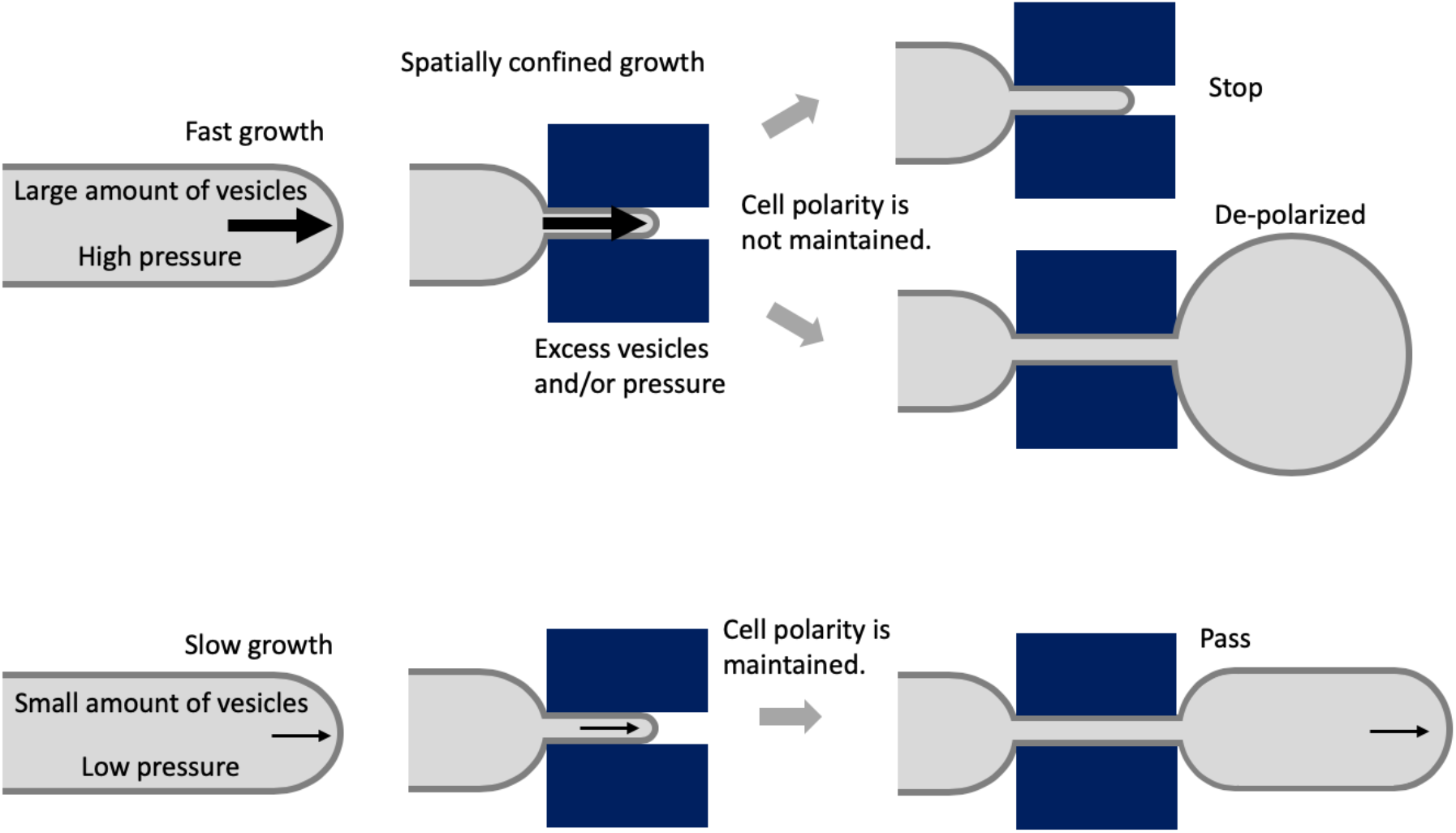
Relationship of growth rate and spatially confined hyphal growth to maintain cell polarity. Cartoon representation of trade-off between cell polarity and growth in spatially confined growth, and how they are correlated and act cooperatively to determine cell shape.

## Discussion

This study showed that hyphae from several fungal species of different phylogenetic lineages were able to grow into microchannels narrower than their width, as described before for plant tip-growing cells (27). It was first found that hyphae of *N. crassa*, *R. oryzae,* and *C. cinerea*, either ceased growing when passing through the channels or became de-polarized upon exiting the channels. The observed effects did not correlate with their taxonomic classification nor with the width of hyphae, but correlated with the hyphal elongation rate. Fast-growing fungi possess the advantage of covering quickly new nutrient-rich substrates or free open spaces. However, at the same time, they may lack the ability to regulate cell shape properly when growing in spatially confined environments. As far as we know this is the first report indicating a trade-off between growth rate and morphogenesis, that suggests the significance of slow growth for the cooperative control of cell polarity and cell growth. This characteristic is considered a case of convergent evolution given the fact that each fungus possesses a similar morphology and physiology adapted to different environmental factors even if they are phylogenetically distant. It will be fascinating in the near future to study whether a similar relationship is observed in other tip-growing cells such as pollen tubes and root hairs of different plant species.

Our results indicate that hyphal tip growth requires a very delicate balance of ordered exocytosis to maintain polarity under spatially constrained circumstances. In fast growing hyphae, such as *N. crassa*, a large number of secretory vesicles are presumably supplied to the hyphal tips, resulting in a conspicuous SPK (43). When fast growing hyphae enter into the narrow channels, a massive number of vesicles is forced to be congregated in the tip region. The cytoplasmic space in those thin hyphae is likely too small for all the secretory vesicles to fit in the tip region. The space constrains probably cause the excess of secretory vesicles to mislocalize at sites others than the tip region, resulting in the de-polarized growth and tip swelling when exiting the channels (Fig. 5). In fact, the lack of localization of some vesicular markers such as CHS-1 at the tip and the dispersed fluorescence observed instead when *N. crassa* hyphae grew through the channels, supports the idea than an excess of vesicles is accumulating in a non-organized manner at the subapical region. When the hyphae are forced to grow through a very narrow channel, under a high turgor pressure, yet maintaining the same growth speed, all the cell-wall building machinery accumulates at the subapical region. Upon exiting the channel, all the machinery gets incorporated in an uncontrolled manner isotropically at the tip, thus generating a swollen tip. After that, new polarity axes get stablished and growth resumes in the form of multiple branches. In the case of *A. nidulans* even if the hyphae are squeezed when entering the channel, the vesicles, presumably less abundant, manage to continue their flow, spacing, movement and grow is non-affected through passage and upon exiting the channel (Fig. 5).

Filamentous fungi play a major role in degradation of biopolymers found in nature for organic material recycling (44, 45). Some fungi are useful in biotechnology and traditional food fermentation (33, 46), where especially solid-state cultivation is important (47). Hyphal invasive growth into the host plant/animal cells is essential for pathogenicity and symbiosis with plant roots as well (38, 48, 49). Our results help to understand the mechanisms of fungal invasive growth into substrates or host cells by spatially confined growth, how cell morphology is controlled by cell polarity and cell growth, that is closely related to fungal biotechnology, ecology and pathogenicity.

## Materials and Methods

### Fungal strains and media

A list of filamentous fungi strains used in this study is given in Table S1. Supplemented minimal medium for *A. nidulans* and standard strain construction procedures are described previously (50).

### Microfluidic device

The microfluidic devices originally designed for culturing tip-growing plant cells and reported by Yanagisawa. et. al (27) were adapted for the current fungal cells studies. Briefly, photoresist (SU-8 3005 & 3010) based microstructures were created on a silicon wafer using a maskless lithography system (DL-1000; Nano System Solutions, Inc.). Then, Polydimethylsiloxane (PDMS, Sylgard 184; Dow Corning) device was prepared through a standard soft-lithography technique. Finally, the PDMS and cover glass (24×60 mm, Matsunami) were both treated with O_2_ plasma (CUTE, Femto Science) for permanent bonding.

### Growth condition

The minimal medium was filled in 20 ml plastic syringe (SS-20ESZ, Terumo) and infused into the PDMS devices using a positive displacement syringe pump (YSP-101, YMC) at a rate of 0.8 μl per hour through a polyethylene tube (Inner diameter 0.38 mm, outer diameter 1.09 mm, BD intramedic).

### Microscopies

Cells were observed by using an epi-fluorescent inverted microscopy, Axio Observer Z1, (Carl Zeiss) microscope equipped with a Plan-Apochromat 63 × 1.4 Oil or 10 or 20 times objective lens, an AxioCam 506 monochrome camera and Colibri.2 LED light (Carl Zeiss). Temperature of the stage was kept at 30°C by a thermo-plate (TOKAI HIT, Japan). Images were collected and analyzed by using the Zen system (Carl Zeiss) and ImageJ software.

### Scanning probe microscope

Cells were grown in the minimal medium on cover slips at 30°C for 24 h. The medium was removed by pipetting then, the cells were analyzed by using a scanning probe microscope SPM-9700HT (Shimadzu) with high magnification optical microscope unit, active vibration isolation table, wide area scanner (XY: 125 μm, Z: 5 μm), and fiber light. Images were collected and analyzed by using Nano 3D mapping software (Shimadzu).

## Acknowledgements

We thank Dr. A. Kogure for the SPM technical assistance, Prof. Y. Kubo for sharing the *C. orbiculare* strain, Prof. H. Muraguchi for sharing the *C. cinerea* strain. This work was supported by the Japan Society for the Promotion of Science KAKENHI Grant (18K05545 to N.T., 19H05364, 20H05412 to Y.S., 18J01077 to N.Y.), Ohsumi Frontier Science Foundation to N.T. and Y.S., and the Japan Science and Technology Agency Exploratory Research for Advanced Technology (ERATO) Grant JPMJER1502.

## Author contributions

N. Takeshita and Y.S. designed research; S.F., R.Y., and N. Takeshita performed research; N.Y., N. Takaya, Y.S. and M.R. contributed new reagents/analytic tools; S.F., R.Y., and N. Takeshita analyzed data; and Y.S., M.R. and N. Takeshita wrote the paper.

---

**Fig. S1.**
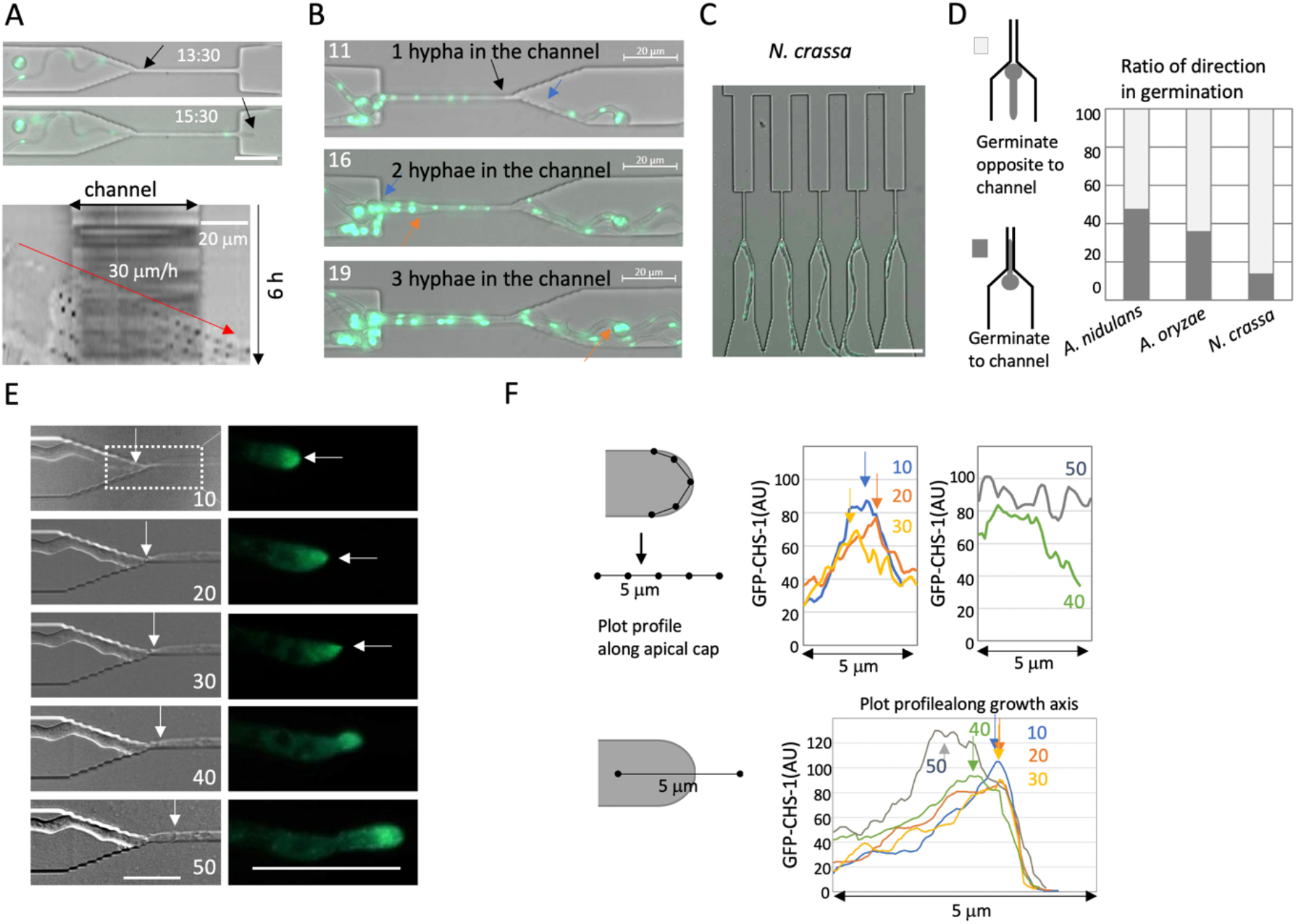
*A. nidulans* but not *N. crassa* hyphae passed through the channels. (A) Time series showing a hypha of *A. nidulans* (nuclei labeled with GFP) growing into the channel. Kymograph along the growth axis before, in and after the channel from Movie S1. The hyphal elongation rates are shown by an arrow. Total 6 h, scale bar: 20 μm. (B) Time series of *A. nidulans* (nuclei labeled with GFP) two or three hyphae passed through the same channel, see Movie S1. Each hyphal tip is shown by arrows. Scale bars: 20 μm. (C) Image of the *N. crassa* spores germinated to the opposite side of slits. (D) Ratio of direction in germination toward or opposite to channels in *A. nidulans, A. oryzae* and *N. crassa*. n=50, respectively. (E) Time series images of *N. crassa* (DIC; left, CHS-1-GFP; right) hyphae growing into a channel from Movie S8. The arrows indicate the SPK. The elapsed time is given in minutes. Scale bar: 20 μm. (F) The plot profile along the apical membrane indicated the signal peaks of SPK (arrows) at 10, 20, 30 min, but not at 40, 50 min. The plot profile along the growth axis (right) indicated the peaks at the apex of hyphae at 10, 20, 30 min, but at the sub-apex at 40, 50 min.

**Fig. S2.**
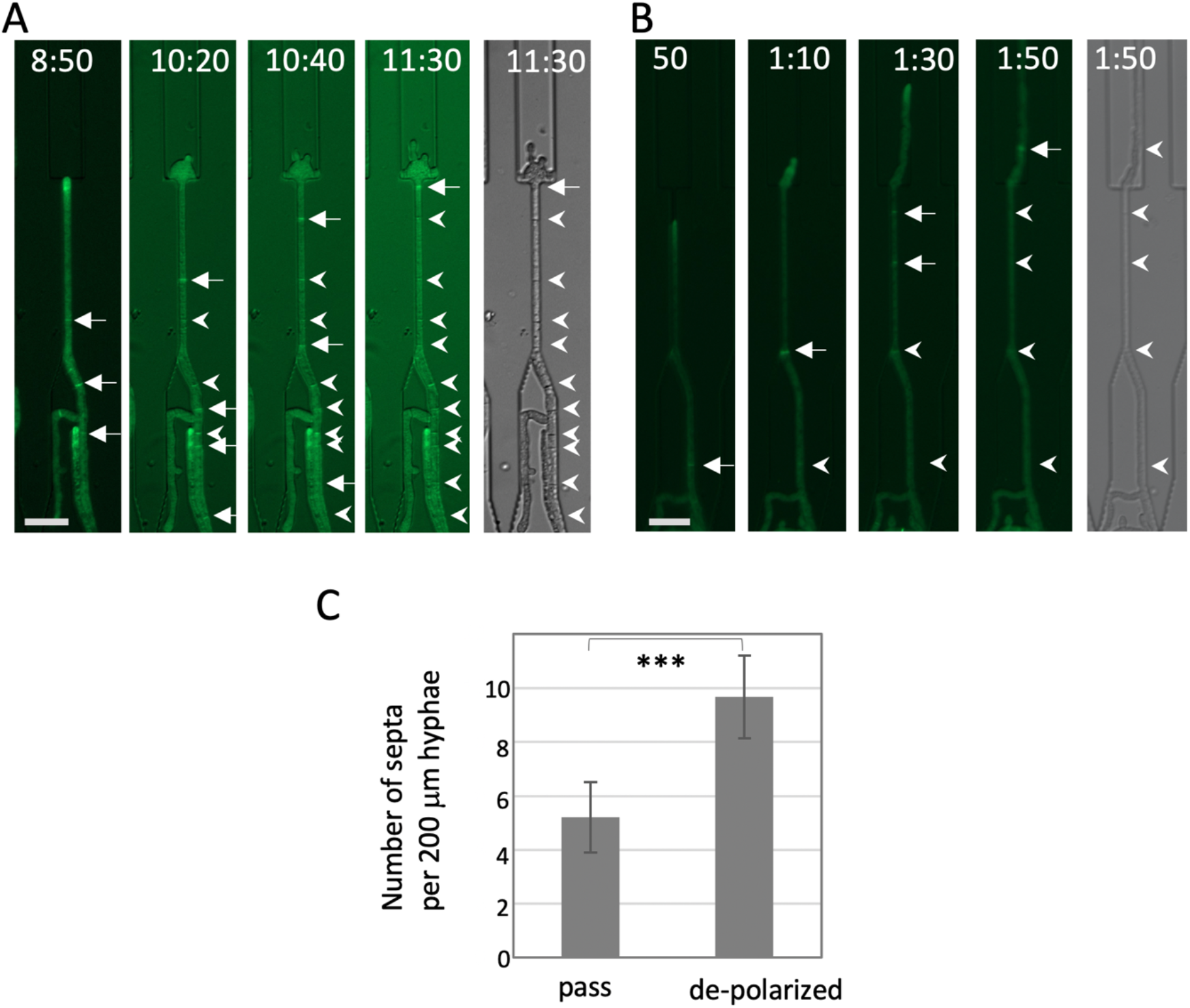
Increased number of septa in de-polarized hyphae. (A, B) Image sequence of forming septa (arrows) and formed septa (arrow heads) in the de-polarized hypha through the channel (A) and in the hypha through the channel (B). The elapsed time is given in hours:minutes. Scale bar: 20 μm. (C) Number of septa in 200 μm-hyphae around the channel, in the hyphae that passed the channels or in the de-polarized hyphae. Error bar: S.D., n = 5, *** P ≤ 0.001.

**Fig. S3.**
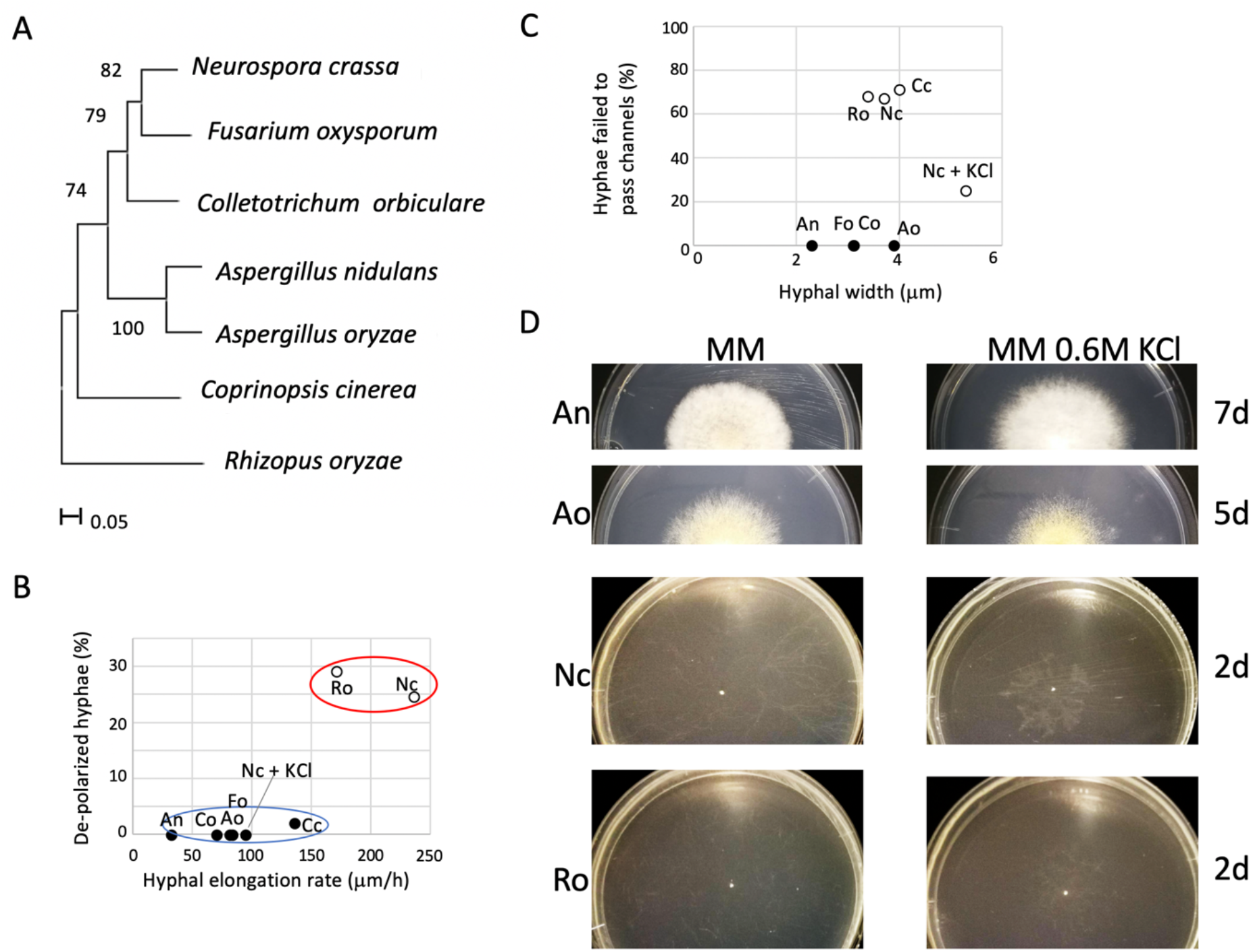
Phylogenetic tree and growth on the plates. (A) Phylogenic tree of filamentous fungi used in this study. Maximum likelihood (ML) tree obtained from the ITS1 and ITS2 regions of the fungal strains. The bootstrap consensus inferred from 100 replicates. (B) Correlation between the hyphal elongation rate and depolarized hyphae. Two groups are shown by red or blue ellipses. (C) No correlation between the hyphal width with the growth defect in channels. (D) Colonies of An; *A. nidulans*, Ao; *A. oryzae*, Nc; *N. crassa* and Ro; *R. oryzae* growth on minimal media (MM) plates or MM + 0.6M KCl plates for 2-7 days.

**Fig. S4.**
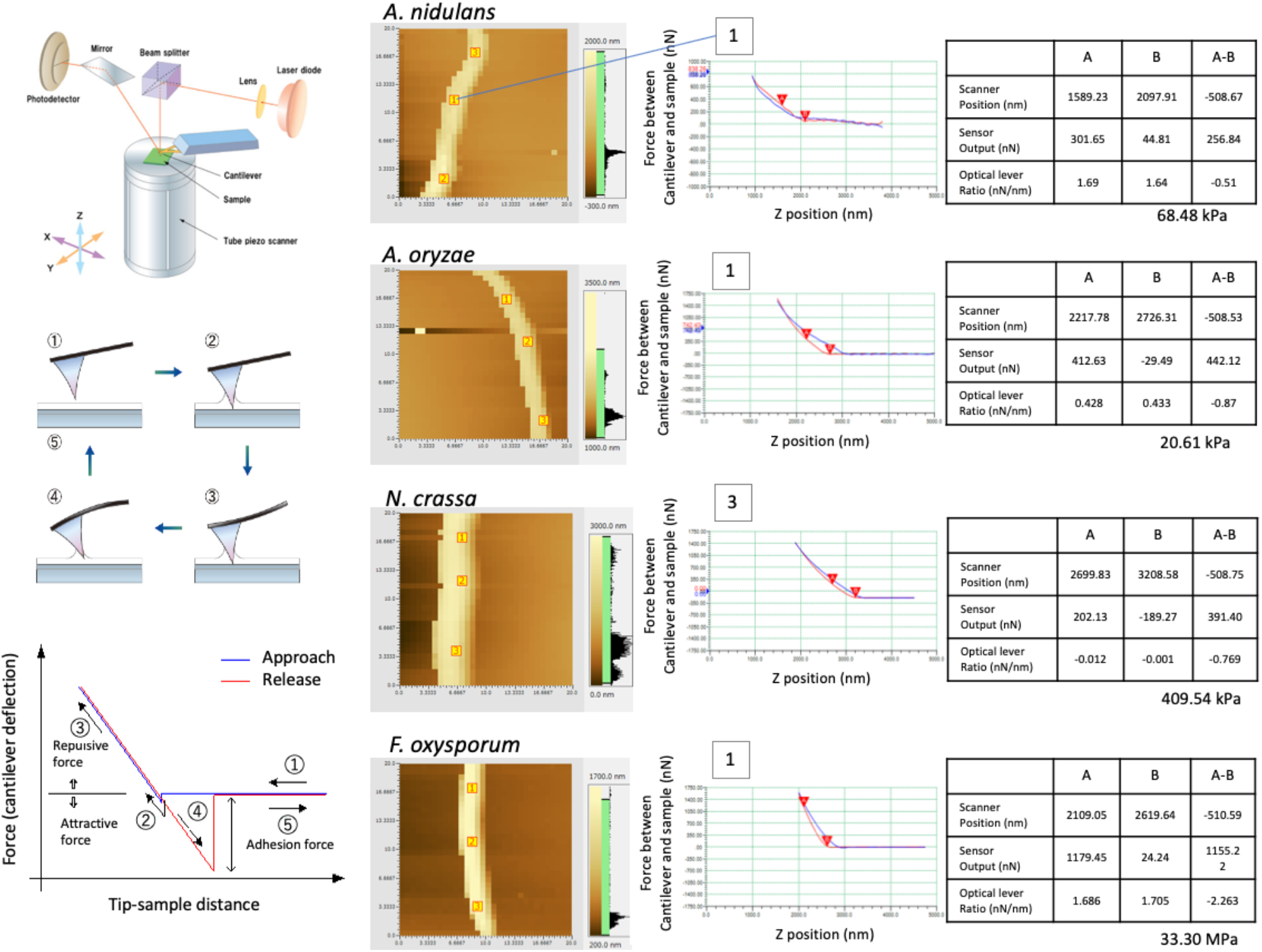
Elastic modules measurement by a scanning probe microscope (SPM). The principle of SPM equipment is composed of the following three. One, laser diode and photo detector. Two, cantilever and holder. Three, scanner*. The basis of the force curve measurement is the measurement performed at one point of the sample. As the distance of the probe changes relative to the sample, this distance can be plotted on the horizontal axis, as shown on the graph. Also, it is possible to calculate from the spring constant of the cantilever and plot this on the vertical axis as nN. When the probe and sample distance is faraway, the force does not work, hence the vertical axis is ①. When the Cantilever touches the sample it is ②. After that, the slope of the graph when the repulsive force acts reflects the hardness of the sample shown as ③. When a release-curve is observed often a large attractive area can be seen. This is because the probe is caught by the adsorption layer on the sample surface shown as ④. From this approach-curve and release-curve, Young’s modulus can be calculated using JKR or Hertz. Therefore, by saving the data at each pixel, a mapping image can be constructed. Fungal cells are grown in a non-invasive manner at high magnifications. *http://www.shimadzu.com/an/surface/spm/faq/index.html

**Fig. S5.**
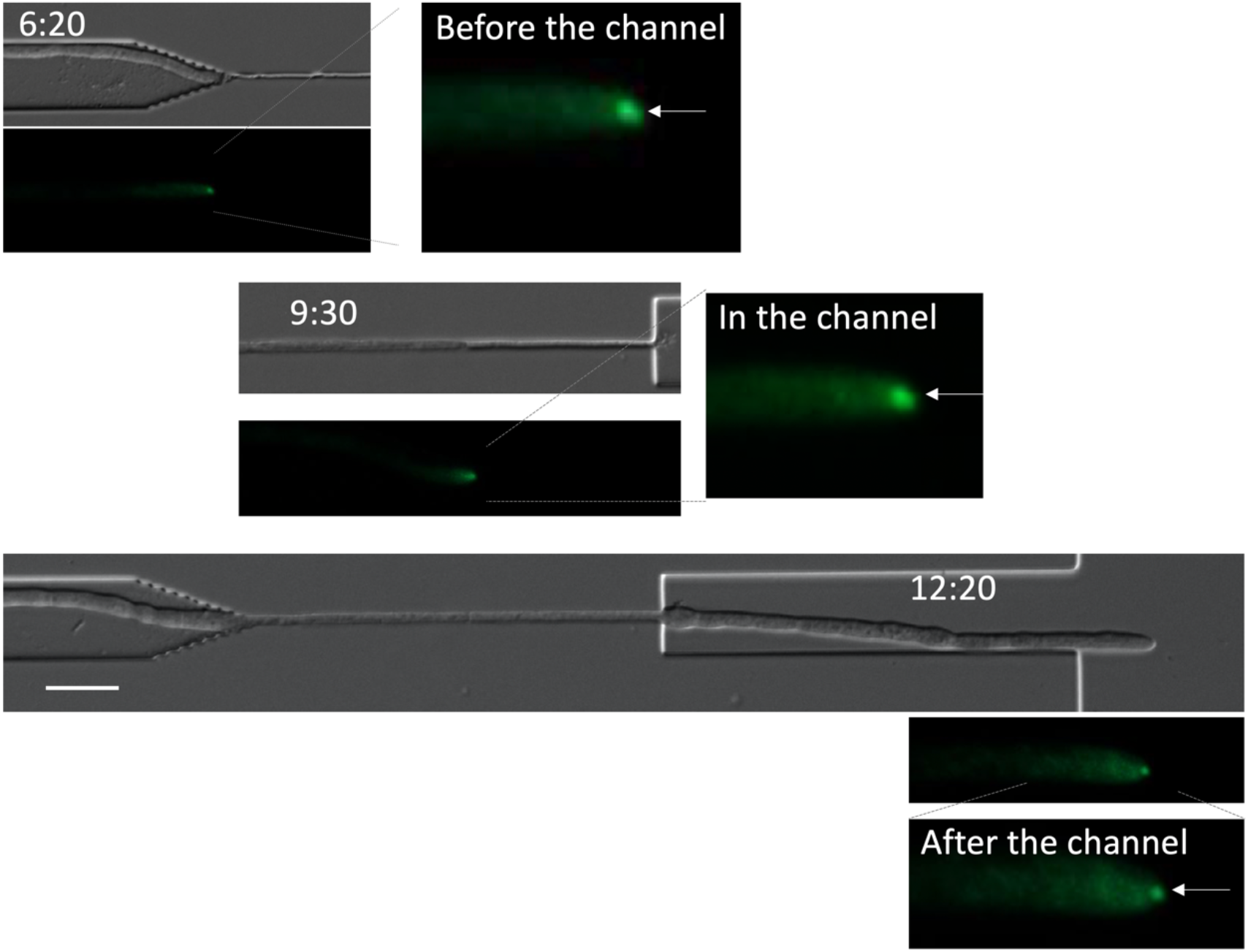
SPK of *N. crassa* hypha grown in MM KCl. Images of the *N. crassa* (SPK labeled with GFP) hypha in the channel from Movie S10. The arrow indicates the SPK before, in and after the channel. The elapsed time is given in hours:minutes. Scale bar: 20 μm.

**Table S1.** Strains used in this study

| Strain | Genotype | Source |
| --- | --- | --- |
| *Aspergillus nidulans* SRS27 | *gpdA* promoter GFP fused StuA-NLS | 1 |
| *Aspergillus oryzae* RIB40UtH2BG | RIB40Δn (pUtH2BG) | 2 |
| *Neurospora crassa* N22813A | *mat A his-3<sup>+</sup>::Pccg-1-hH1<sup>+</sup>-sgfp<sup>+</sup>* | 3 |
| *Neurospora crassa* NES1-15 | *mat A his-3<sup>+</sup>::Pccg-1::chs-1::sgfp<sup>+</sup>* | 4 |
| *Fusarium oxysporum* JCM11502 | Wild type | JCM: Japan Collection of Microorganisms |
| *Colletotrichum orbiculare* 104-T | Histone H1-GFP | 5 |
| *Rhizopus oryzae* JCM5582 | Wild type | JCM |
| *Coprinopsis cinerea* #2 + #8 dikaryon | A3B1 + A2B2 | 6, 7 |

## Movie Legends

Movie S1. *Aspergillus nidulans* strain whose nuclei were visualized by GFP grew into, through and out of the channels. Every 10 min, total 10h, scale bar: 20 μm.

Movie S2. *Aspergillus oryzae* strain whose nuclei were visualized by GFP grew into, through and out of the channels. Every 20 min, total 14h, scale bar: 50 μm.

Movie S3. *Neurospora crassa* strain whose nuclei were visualized by GFP often showed de-polarized hyphae out of the channels. Every 5 min, total 20 h, scale bar: 20 or 50 μm.

Movie S3. *Neurospora crassa* strain whose nuclei were visualized by GFP often showed de-polarized hyphae out of the channels. Every 5 min, total 20 h, scale bar: 20 or 50 μm.

Movie S4. *Neurospora crassa* strain expressing GFP-CHS-1 penetrated into the channels then stopped growing. Every 10 min, total 50 min, scale bar: 20 μm.

Movie S5. *Neurospora crassa* strain expressing GFP-CHS-1 penetrated into the channels then showed de-polarized hyphae out of the channels. Every 10 min, total 6 h, scale bar: 20 μm.

Movie S6. *Fusarium oxysporum* strain grew into, through and out of the channels. Every 20 min, total 7 h, scale bar: 50 μm.

Movie S7. *Colletotrichum orbiculare* strain grew into, through and out of the channels. Every 20 min, total 16 h, scale bar: 50 μm.

Movie S8. *Rhizopus oryzae* strain showed de-polarized hyphae out of the channels. Every 10 min, total 15 h, scale bar: 50 μm.

Movie S9. *Coprinopsis cinerea* dikaryon strain penetrated into the channels then stopped growing. Every 10 min, total 24 h, scale bar: 50 μm.

Movie S10. *Neurospora crassa* strain expressing GFP-CHS-1 grew into, through and out of the channels in the high osmotic condition. Every 10 min, total 15 h, scale bar: 50 μm.

